# Computational-Observer Analysis of Illumination Discrimination

**DOI:** 10.1101/302315

**Authors:** Xiaomao Ding, Ana Radonjić, Nicolas P. Cottaris, Haomiao Jiang, Brian A. Wandell, David H. Brainard

## Abstract

The spectral properties of the ambient illumination provide useful information about time of day and weather. We study the perceptual representation of illumination by analyzing measurements of how well people discriminate between illuminations across scene configurations. More specifically, we compare human performance to a computational-observer analysis that evaluates the information available in the isomerizations of the cones in a model human photoreceptor mosaic. Some patterns of human performance are predicted by the computational observer, other aspects are not. The analysis clarifies which aspects of performance require additional explanation in terms of the action of visual mechanisms beyond the isomerization of light by the cones.

## Introduction

The spectral properties of the ambient illumination provide useful information about time of day and weather. Indeed, variation in illumination spectra occurs outdoors over the course of the day (e.g., Hernandez-Andres, Romero, Nieves, & Lee, 2001; Spitschan, Aguirre, Brainard, & Sweeney, 2016), within single scenes (Nascimento, Amano, & Foster, 2016), and when we move between natural and artificial illumination (Wyszecki & Stiles, 1982). A number of psychophysical paradigms have been developed to study the perceptual representation of illumination (Kardos, 1928; Beck, 1959; Oyama, 1968; Kozaki & Noguchi, 1976; Gilchrist & Jacobsen, 1984; Noguchi & Kozaki, 1985; Hurlbert, 1989; Logvinenko & Menshikova, 1994; Rutherford & Brainard, 2002; Logvinenko & Maloney, 2006; Lee & Brainard, 2011). Recently, a number of labs have reported psychophysical threshold measurements of how well people can discriminate between illuminations across scene configurations (Pearce, Crichton, Mackiewicz, Finlayson, & Hurlbert, 2014; Radonjić et al., 2016; Alvaro, Linhares, Moreira, Lillo, & Nascimento, 2017; Radonjić et al., 2018).

A key step in interpreting psychophysical threshold measurements is to understand the degree to which patterns in the data are driven by variation in the information available from the stimuli. This has often been accomplished by comparing human performance to that of an *ideal observer* that makes optimal use of the task-relevant information available in the stimulus or at some early stage of the visual processing (Barlow, 1962; Green & Swets, 1966; Geisler, 1989; Geisler, 2011). Such ideal observer analyses clarify which aspects of performance may be accounted for by the properties of the stimuli and well-understood mechanisms of early vision. The optical point spread function of the eye, for example, accounts for much but not all of the decrease in human spatial contrast sensitivity at high spatial frequencies (Banks, Geisler, & Bennett, 1987; Sekiguchi, Williams, & Brainard, 1993).

In this paper, we review previously reported measurements of human psychophysical performance on an illumination discrimination task (Radonjić et al., 2016; Radonjić et al., 2018). We then compare human performance to that of a computational observer that uses the information available in the cone photoreceptor isomerizations to perform the same task. We ask whether the stimulus dependent changes in human performance across different illumination changes and scene configurations are consequences of differences in the information available to perform the task. The analysis clarifies which aspects of performance require additional explanation in terms of the action of visual mechanisms beyond the isomerization of light by the cones.

Our approach shares much with the ideal observer analysis, but rather than using an analytic calculation to estimate ideal performance levels, we employ computer simulations and machine learning. For this reason, we refer to our approach as a *computational observer analysis* (Farrell, Jiang, Winawer, Brainard, & Wandell, 2014; Jiang et al., 2017; cf. Lopez, Murray, & Goodenough, 1992). Unlike the ideal observer, where the decision rule is implemented on the assumption that the observer has perfect information about the statistical properties of the stimuli, the computational observer must learn these properties from a finite number of training samples.

## Illumination Discrimination Psychophysics: Modeled Experiment 1

We analyze two similar illumination discrimination experiments; we begin by providing an overview of the first. Detailed methods and results for this experiment (Modeled Experiment 1) are reported elsewhere (Radonjić et al., 2018, Experiment 2 in that paper, fixed-surfaces condition), so our overview is brief. The second experiment (Modeled Experiment 2) is similar in design and is also reported in detail elsewhere (Radonjić et al., 2016).

Both modeled experiments measured human’s ability to discriminate changes in illumination across four illumination-change directions – blue and yellow (which were aligned with the daylight locus) and red and green (which were orthogonal to the daylight locus). On each trial of the experiments, observers viewed three successive computer-generated images, displayed on a calibrated color monitor. The scene geometry was held fixed across the experiment and all the surfaces in the scene remained unchanged: only the spectral power distribution of the illumination varied. The first image on a trial was presented in the *reference interval* (2370 ms). For this image, the scene was illuminated by the *target illumination*, a metamer for natural daylight with a color temperature of approximately 6700 K. The reference interval was followed by two *comparison intervals.* The comparison intervals (870 ms) were separated from each other and from the reference interval by inter-stimulus intervals (750 ms). During each comparison interval, an image of the scene was again shown. In one comparison interval, the scene was illuminated by the target illumination and in the other by a *test illumination.* The order of the two comparisons was randomized on each trial. The observer’s task was to choose which of the two comparison intervals had scene illumination most similar to the target illumination. On each trial, the test illumination was chosen from a pool of 200 pre-specified illuminations. These varied in steps of approximately 1 CIELUV ΔE unit (CIE, 2004) in each of four illumination-change directions: blue, yellow, green and red (50 steps per direction).^1^ Figure 1 shows example images of the scene under the target illumination and four test illuminations.

**Figure 1.**
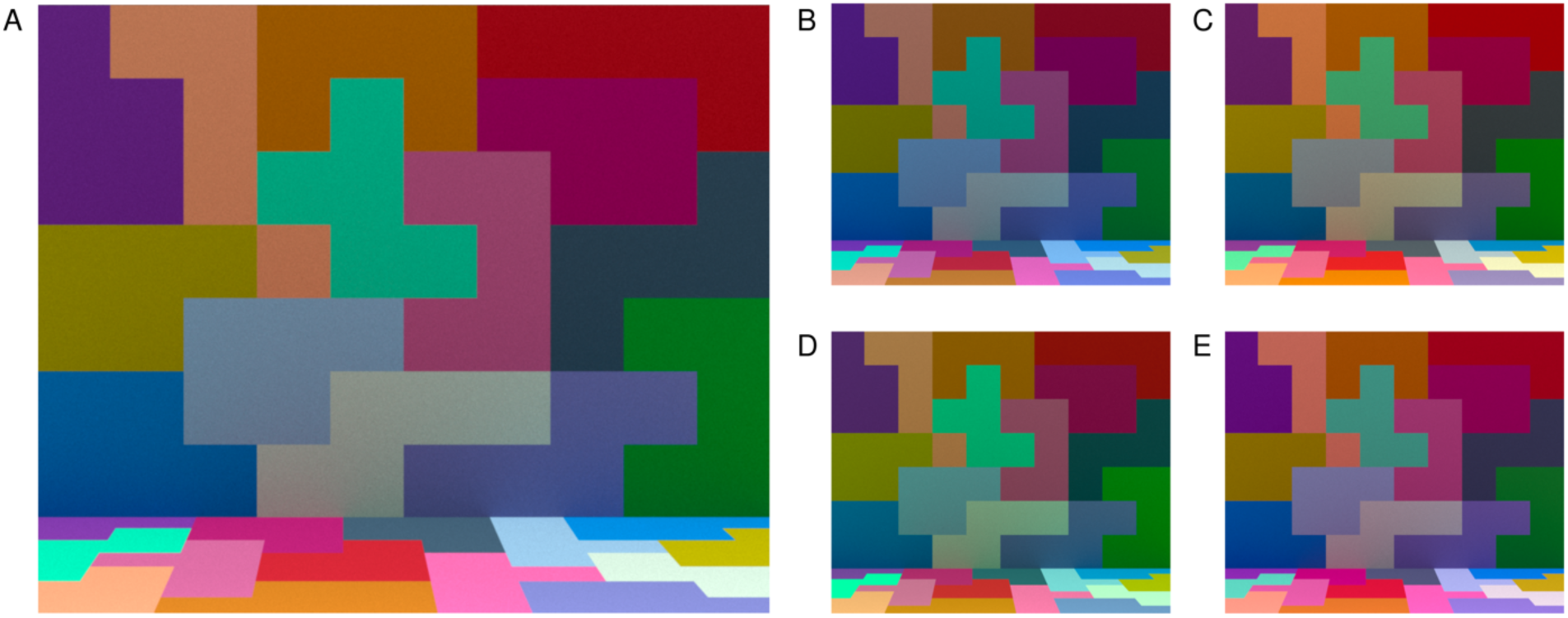
Experimental stimuli. **A)** Image of the scene illuminated by the target illumination. **B-E)** Images of the scene illuminated by four test illuminations. Each illumination is approximately 30 ΔE steps away from the target. Panels B-D correspond to test illuminations in the blue, yellow, green, and red illumination-change directions respectively. Images in B-E are shown at a smaller size than in A to conserve space. In the experiment, the target and test illumination images were the same size. Images rendered here and in other figures in this paper are illustrative. The rendering process is unlikely to preserve the precise color appearance of the experimental stimuli.

During each session, trials probing the four illumination-change directions were interleaved. The illumination-change steps were determined through twelve interleaved 1-up-2-down staircases (three independent staircases for each illumination-change direction). The illumination step size for a staircase was decreased when the observer responded correctly twice in a row on trials governed by that staircase, and increased each time the observer responded incorrectly. Each staircase terminated after the 8^th^ reversal.

Observers’ thresholds to discriminate illumination changes were measured for each of the four directions, relative to the target illumination. Thresholds corresponded to 70.71% accuracy and were determined using a maximum likelihood fit to the combined data for all three staircases in each illumination-change direction. The 70.71% correct criterion was used because this is the performance convergence point for the staircases (Wetherill & Levitt, 1965). Observers completed two experimental sessions, and thresholds obtained in each session were averaged for each observer. Each observer’s right eye was tracked using an Eyelink 1000 (SR Research, Ltd.), which allowed us to record eye fixation positions on each trial.

Figure 2 illustrates the average thresholds across ten observers. The average threshold for the blue direction is elevated relative to those obtained for the other illumination-change directions, when thresholds are plotted using the CIELUV ΔE metric. The elevation for thresholds in the blue direction has been found in other studies with a similar design, for scenes where the average reflectance of the surfaces in the scene is close to that of a neutral nonselective gray (Pearce, Crichton, Mackiewicz, Finlayson, & Hurlbert, 2014; Radonjić et al., 2016). Two essentially identical experiments reported with this one (Radonjić et al., 2018, Experiments 1 and A1 in that paper, fixed-surfaces condition) yielded similar results. Here we ask We are particularly interested here in whether the elevation of threshold in the blue direction has its roots in a relative paucity of information about this illumination change in the representation of the visual scene provided by the cone photoreceptor mosaic.

**Figure 2.**
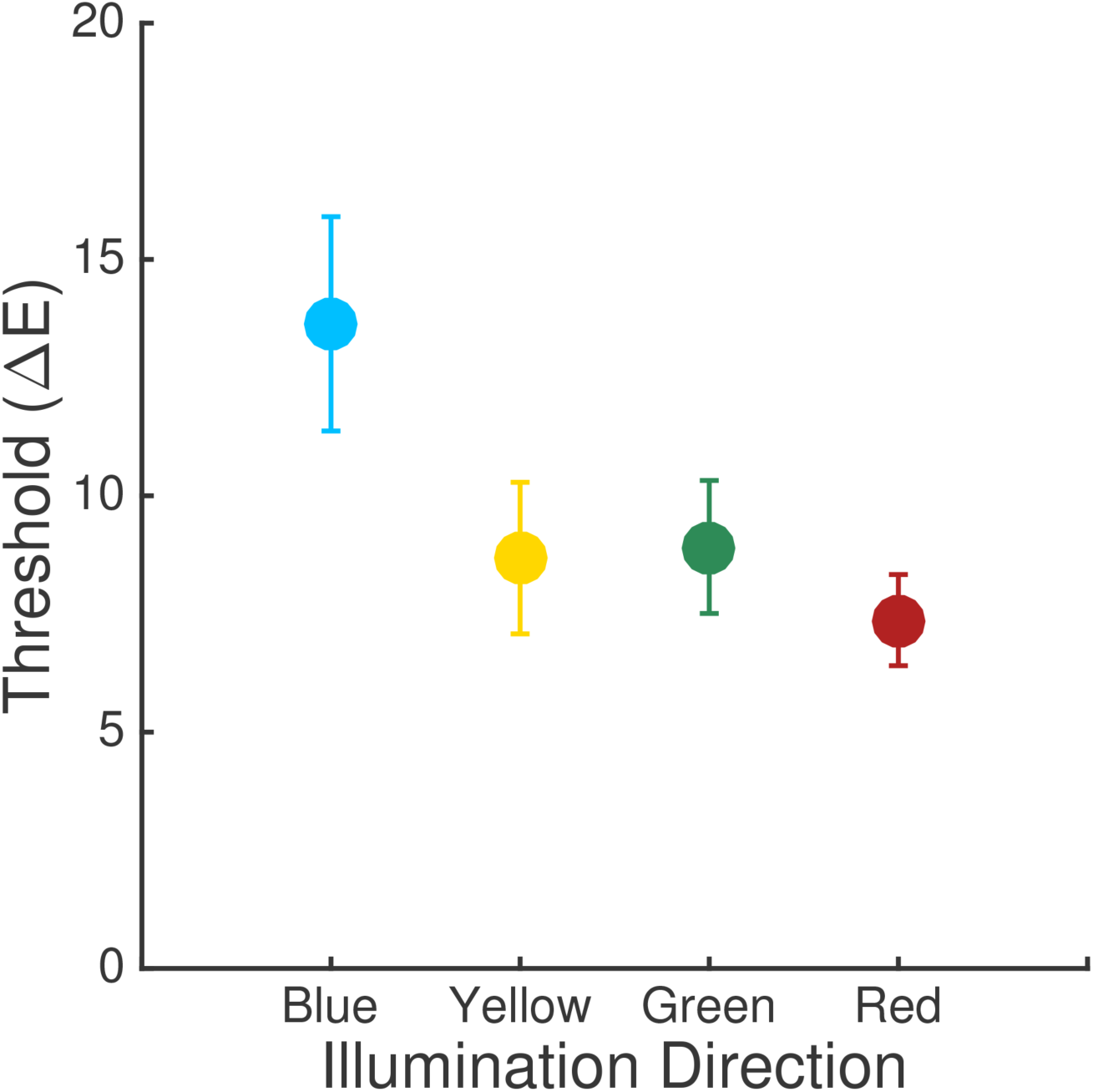
Experimental results. Thresholds averaged across observers for Experiment 1. Error bars are +/−1 SEM. Each point shows the threshold for one illumination-change direction.

## Computational-Observer Analysis

We analyze the information available for performing the illumination discrimination task that is carried by the isomerization rates of the cone photoreceptors. We used the Image System Engineering Toolbox for Biology (ISETBio; isetbio.org) to model retinal image formation and the absorption of light in the foveal cone mosaic. ISETBio provides functions that model early vision and allow computation of the visual system’s responses to calibrated stimuli at various stages of processing. It includes routines for computation of the retinal image from a description of scene radiance (including blurring by the eye’s optics and absorption of light by the lens and macular pigment), as well as computation of the photopigment isomerizations of each cone in a patch of photoreceptor mosaic (while taking into account the spectral sensitivities of the three classes of cones in the human retina and their interleaved spatial sampling). ISETBio also contains routines for learning observer decision rules from labeled training data, and we employ these routines to determine the performance of our computational observer.

ISETBio is written in Matlab (The MathWorks, Natick, MA) and is freely available under an open-source software license (isetbio.org). The code and data required to use ISETBio to reproduce the calculations and figures in this paper are available at https://github.com/isetbio/BLIlluminationDiscriminationCalcs.

### Modeling of the Stimuli

The stimuli were presented on calibrated computer monitors with known size and distance from the observer. In Experiment 1, stimuli were presented to the right eye only and we modeled the information contained in the right eye image. In Experiment 2, the stimuli were presented stereoscopically, but for simplicity we modeled only the information contained in the left eye image. We imported the RGB stimulus images into the ISETBio *scene* format, which uses the display calibration information to compute the spectral power distribution of the light emitted from each location in the image. The scene representation also specifies the size of the image and the distance between the display and the eye. These values were set to approximate their experimental values.^2^

### The Retinal Image

We used ISETBio routines to compute the retinal image from each stimulus image. In ISETBio terminology, this is referred to as the *optical image.* These routines incorporate the size of the pupil (set to 3 mm in our calculations), the geometry of image formation, absorption of light by the lens and blurring by the eye’s optics. We used the ISETBio default estimates of lens density from Bone, Landrum, and Cairns (1992) and the polychromatic shift-invariant model of the optics of the human eye provided by Marimont and Wandell (1994).

### Cone Isomerizations

We estimated the mean number of isomerizations in the cones for a 1° × 1° patch of foveal retina. We chose a 1° × 1° patch of retina because restricting the analysis to patches of this size makes calculation of computational-observer performance tractable given current computing power. The full analysis is thus achieved by breaking the image down into a set of 1° × 1° patches and aggregating performance over these patches.

The model cone mosaic was rectangularly packed with an inter-cone spacing of 2 um. This corresponds to the peak foveal density reported by Curcio et al. (1991). The class of each cone (L, M or S) at each location was chosen randomly. There were 22201 cones in all, 13765 (62%) L cones, 6882 (31%) M cones and 1554 (7%) S cones, corresponding to current views on the relative numbers of cones of each class in the typical human retina (Hofer, Carroll, Neitz, Neitz, & Williams, 2005). Photopigment absorbance for the L, M and S cones was taken from the CIE 2007 standard (CIE, 2007), and absorption by the foveal macular pigment (Stockman, Sharpe, & Fach, 1999) was incorporated. These choices, together with that used for lens density above, yield the CIE (2007) 2° cone fundamentals. We assumed a quantal efficiency for cone photopigment of 0.67 (Rodieck, 1998), a foveal cone inner segment acceptance area of 1.96 um^2^, an area that sits between those provided by Kolb (http://webvision.med.utah.edu) and Rodieck (1998). We used a cone integration time of 50 ms, which allowed us to compute the mean number of isomerizations of each cone in response to each image. From the mean number of isomerizations, we can simulate the number of isomerizations on an individual trial, because trial-by-trial isomerizations are Poisson distributed (Hecht, Schlaer, & Pirenne, 1942; Rodieck, 1998).

### Obtaining Computational-Observer Thresholds

Given the responses of a simulated cone mosaic to the experimental stimuli, we made
computational-observer predictions of discrimination performance for each illumination-change direction and step size. To do so, we created a training set consisting of 1000 simulated trials and used this to learn a linear classifier that separated trials according to which comparison interval corresponded to the target. Each instance in the training set had independently drawn noise added to the mean number of isomerizations for each cone. We predicted performance by evaluating the classifier on a separate test set (cross-validation). The test set consisted of an additional 1000 simulated trials, each with independently drawn noise added.

We studied performance for a 1° by 1° patch of foveal mosaic and a single fixation location, which was held constant across the two comparison intervals. This quantifies the information available from a single patch of the stimulus. We then investigated how this information varies across stimulus patches and related this to where observers look during the experiment. Our approach ignores the additional information provided in the reference interval. The motivation for including that interval in the experiment was to reduce observer’s uncertainty about which of the two comparison-interval images corresponds to the target. Our computational observer was designed so that such uncertainty is not an issue.

For each illumination-change direction and step size, we calculated the mean number of isomerizations of the cones in the mosaic to the target-illumination image patch and the test-illumination image patch. We used an 50 msec cone integration time. To each mean response, we added Poisson noise to simulate trial-to-trial variability in the number of isomerizations in a single integration time. Then, we concatenated the resulting two response vectors into a single vector, with the target coming either first or second. We refer to simulated trials where the target vector comes first and the comparison vector second as AB trials, and simulated trials where the comparison vector comes first and the target second as BA trials. The task of the computational observer was to classify AB versus BA response vectors.

The training and test sets each consisted of 1000 labeled concatenated response vectors of this sort (500 AB and 500 BA). We used the training set to learn a linear classifier, i.e., to find the hyperplane which best separated the two classes (AB versus BA) of concatenated response vectors. We did this using the support vector machine (SVM) algorithm (Manning, Raghavean, & Schutze, 2008), as implemented in Matlab (function fitcsvm). This algorithm finds the hyperplane that maximizes the margin between the exemplars of the two classes, where margin refers to the amount of space without any data points around the classification boundary. Support vector machines provide an effective tradeoff between good classification performance on the training set and good generalization.

The dimensionality of the concatenated response vectors is 44402 for two concatenated 1° by 1° patches of foveal retina. We reduced this dimensionality using principal components analysis (PCA). We first standardized each dimension (i.e., response of one cone in one interval) of the training set using its sample mean and standard deviation. Then, we ran principal component analysis on the training set and projected the standardized response vectors onto the first 400 principal components for training and testing.

Figure 3A shows the projection of 100 AB and 100 BA vectors from one stimulus patch, using nominal step size ΔE = 1 in the blue illumination-change direction, onto the first 2 principal components obtained for this direction and step size. The dashed line shows the decision boundary of a linear SVM trained on these vectors. Using this decision boundary led to performance of 100% on the training set as well as on the test set, indicating that a computational observer could perform perfectly on the task even for this smallest illumination-change step size. Since human thresholds considerably exceed ΔE = 1, there are additional sources of noise beyond the Poisson variation in the number of isomerizations and/or inefficiencies in the way that the visual system uses the information provided by the cone isomerizations. Indeed, ideal observer studies of human discrimination performance for simple stimuli often find that ideal observers outperform human observers (Geisler, 1989). Since our goal is to understand whether the pattern of psychophysical performance we measured is driven by differences in information at the photoreceptor mosaic, we modeled the inefficiency of the post-receptoral visual system by adding independent zero-mean Gaussian noise to each cone’s response, in addition to the Poisson noise. Doing so degrades the performance of the computational observer, but in a manner that does not seem likely to introduce systematic changes in relative computational-observer performance across illumination-change directions. We set the variance of the added noise for each cone’s response (both for the target and comparison responses) to a multiple of the mean response of the cones in the mosaic; we refer to the multiple chosen as the *noise factor.* The noise factor was varied systematically, providing us with a parameter that allowed us to match performance levels between computational and human observers.

**Figure 3.**
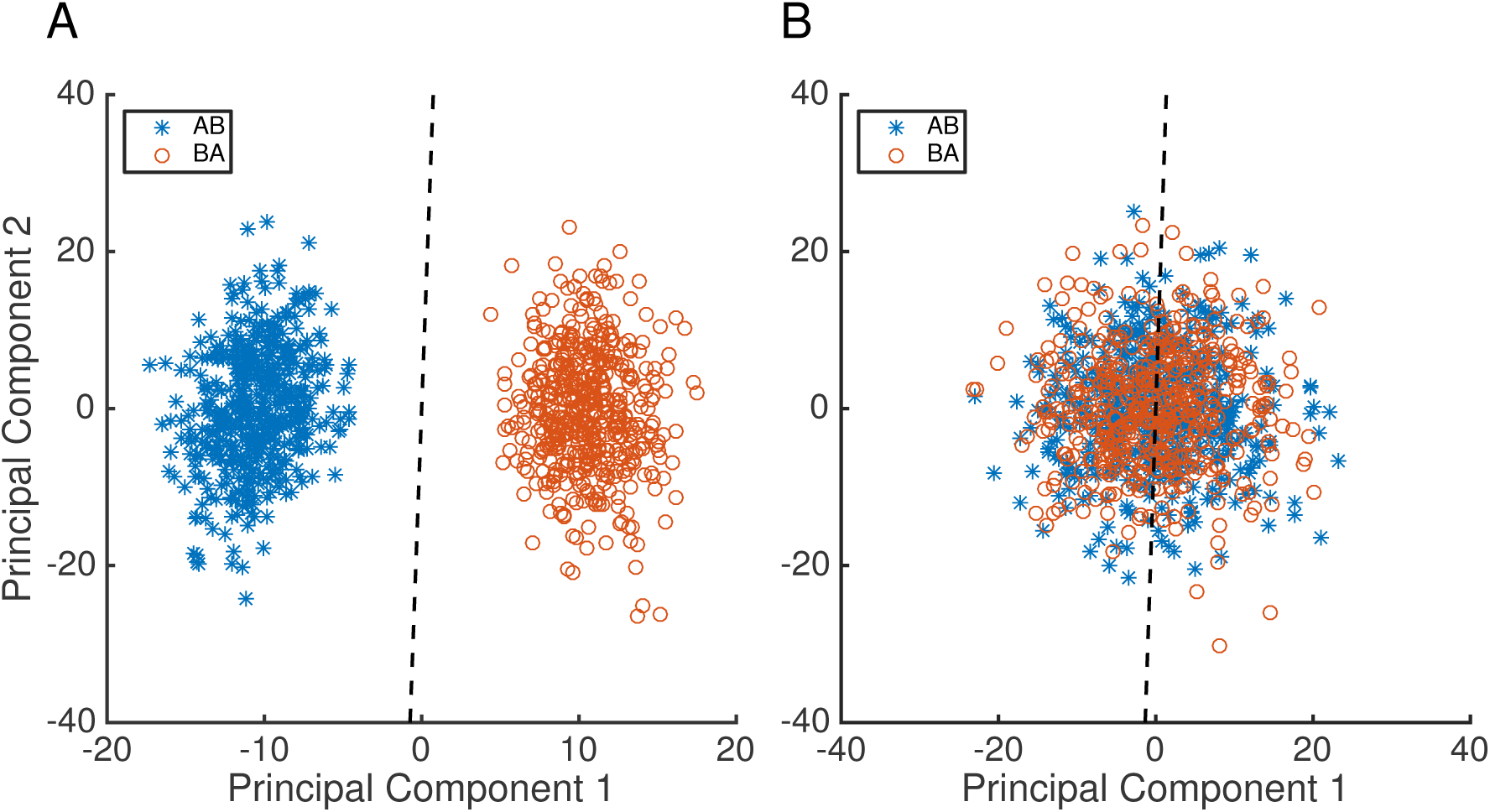
Examples of learned classification boundaries. **A)** Plot of 100 samples of AB data vectors and 100 samples BA data vectors projected onto the first two principal components of the cone mosaic response space. Data are for the illumination-change step size (ΔE = 1) in the blue direction. The dashed line is the linear SVM decision boundary learned from the 200 samples in this training set. The variability is due to Poisson noise; no Gaussian noise was added. **B)** Corresponding plot with 15× additive Gaussian noise added to the responses, prior to data reduction by PCA and classification learning. Note that the principal components are not identical between the two panels, as the PCA calculation was done separately on the two training sets.

Figure 3B shows the same 100 AB and BA vectors from 3A, but with 15× (noise factor) Gaussian noise added to the original responses. These noisy responses are projected onto principal components computed from a training set of response vectors with the same noise factor. The two classes are now overlapping, illustrating how adding the Gaussian noise reduces performance. Indeed performance on the test case for this set is essentially at chance (51%)

To model the psychophysical data, we trained SVMs for each combination of illumination direction, step size and a series of noise factors. For each combination, we applied principal components analysis to the standardized training set (1000 instances), projected each instance onto the first 400 dimensions, and then trained the SVM. To evaluate performance for the combination, we generated an additional 1000 concatenated response vectors to form a test set, standardized these using the means and variance from the training set, projected these onto the 400 dimensions of the PCA for the training set, and classified using the SVM. This provides us with a modeled percent correct for each combination. Noise factors were varied from 0 to 30 in steps of 5.^3^

As noted above, it was not computationally feasible for us to train the SVM’s using a cone mosaic that captures the entire stimulus. For this reason, we partitioned the scene into 1° × 1° square patches by imposing a uniform grid onto the stimulus, and then trained SVM’s for each patch across all combinations of illumination direction, step size, and noise factor. Figure 4A illustrates the grid. On the right and bottom edges of the image, there are small regions of the stimulus not covered by the grid. These are neglected in the calculations. To obtain threshold for each illumination-change direction and choice of noise factor for each 1° × 1° patch, we analyze computational-observer performance as a function of step size by fitting a Weibull psychometric function. This relates computational-observer performance to illuminant-change step size. In performing the fit, the actual ΔE values for each illumination change, rather than the nominal values, were used (see Radonjić et al., 2018). The fit was obtained using the maximum-likelihood method implemented in the Palemedes Toolbox (Kingdom & Prins, 2010; www.palamedestoolbox.org, version 1.8.2). Figure 4B shows computational-observer performance and the fit, for the blue illumination-change direction at 10× noise factor, for the patch outlined in white in the upper left (3^rd^ row from the top, 1^st^ column from the left) of Figure 4A. Using the fitted psychometric functions, 70.71% correct thresholds were extracted.

**Figure 4.**
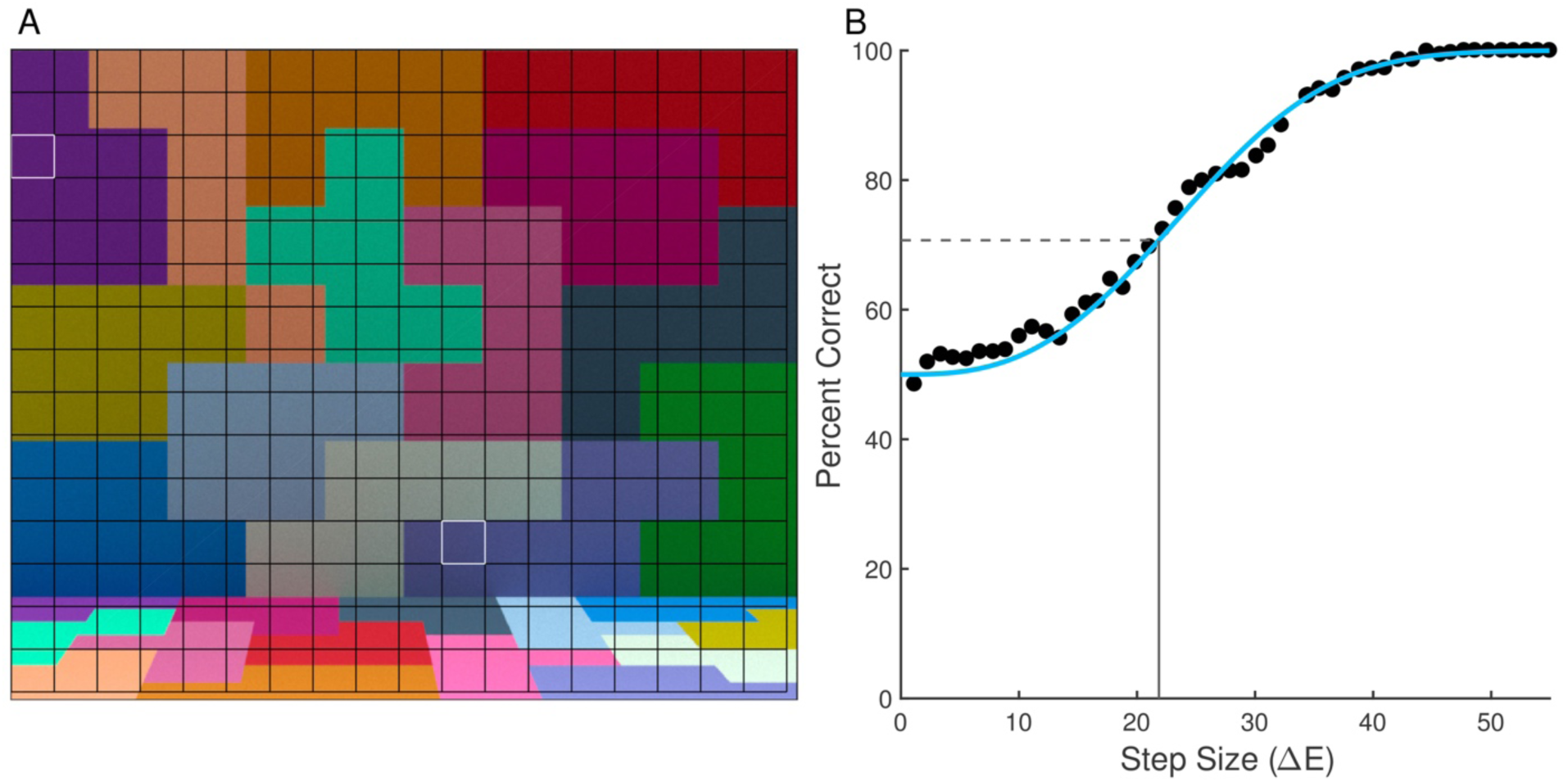
Obtaining computational-observer thresholds. **A)** Stimulus with uniform grid superimposed on top. Each square in the grid is 1° × 1° visual angle. Each image square was analyzed separately. **B)** Plot of linear SVM performance at each stimulus level for one stimulus patch (row 3, column 1); white outline in A). Points represent the performance of a linear SVM trained and tested for each step size for the blue direction with 10× additive Gaussian noise factor. The bestfit Weibull psychometric function is depicted with the solid blue line. The horizontal dashed grey line is the 70.71% accuracy level and the vertical solid grey line is the threshold (22 ΔE units in this example), calculated using the inverse of the fit function. Performance for the patch in outlined in white in the lower center of 4A (row 12, column 11) is shown in Figure 5B.

Figure 5A shows the computational-observer thresholds for each illumination direction plotted as a function of noise factor, for the same stimulus patch (row 3, column 1) whose performance is shown in Figure 4B. For low added noise, thresholds are very low, as one would expect based on Figure 3A. As the noise factor increases, thresholds also increase. The relative ordering of the thresholds across illumination-change directions is preserved across noise factors (yellow > blue > green > red).

**Figure 5.**
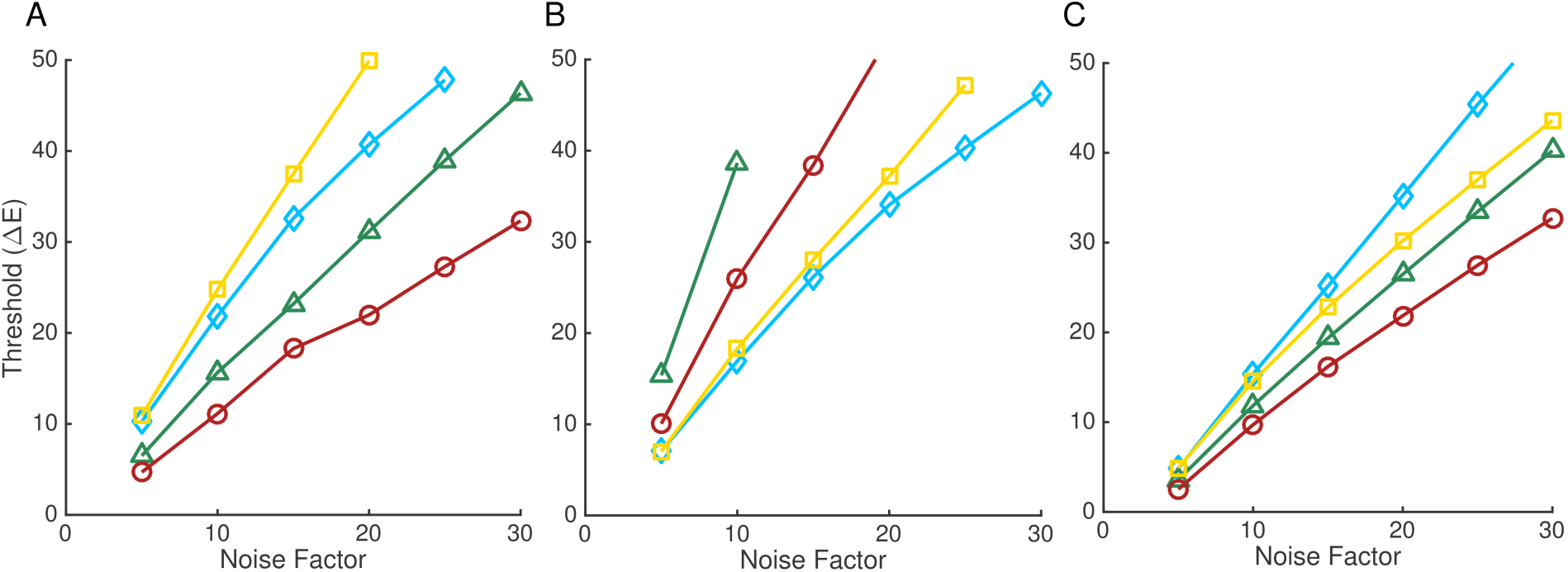
Computational-observer thresholds as a function of noise factor. **A)** Computational-observer thresholds from a single stimulus patch (top left white outline in Figure 4A) for the four illumination-change directions (blue: blue diamonds; yellow: yellow squares; green: green triangles; red: red circles), as a function of noise factor. **B)** Same as A but for a different stimulus patch (white outline in the lower-center portion of Figure 4A). The line for the green illumination-change direction does not extend further because at higher noise levels performance was too poor to estimate a threshold. **C)** Mean thresholds as a function of noise factor, obtained by averaging performance over all the patches shown in 4A for each illumination direction/step size/noise factor. The psychometric function is fit to the average values.

Figure 5B shows performance for a different stimulus patch, indicated by a white outline in the lower-center portion of Figure 4A (row 12, column 11). Thresholds again increase with noise factor, but here the ordering of thresholds in the different illumination-change directions is different than in 5A (green > red > yellow > blue). This effect is due to the difference in surface reflectance at the two patches. The two patches for which performance is illustrated in Figure 5A and B were chosen to illustrate the large effect that patch choice can have on computational-observer performance.

Because predicted performance depends on which patch is examined, to make overall predictions it is necessary to aggregate across the individual patches. There are a number of ways to implement such aggregation. Here, for each illumination direction, step size and noise level, we averaged the percent correct performance obtained for each of the 270 patches (Figure 4A) to obtain a single aggregate psychometric function. We then analyzed this function using the same methods as for the individual patch data, to obtain an aggregate threshold as a function of noise factor. The results of this analysis are shown in Figure 5C. The aggregate performance more closely resembles the performance in 5A than in 5B, but differs in relative threshold order (blue > yellow > green > red). The aggregate thresholds represent an estimate of computational-observer performance when the information in all patches is weighted equally.

### Relation to Psychophysics

For each human observer, we found a single noise factor that minimized the sum of the squared error (across the four illumination-change direction) between the aggregate computational-observer thresholds (Figure 5C) and human thresholds. Linear interpolation was used to estimate computational-observer thresholds for noise factors between those for which performance was explicitly computed. The resulting computational-observer thresholds across observers were averaged to provide a fit to the average human data. This average fit is shown in Figure 6A. The computational-observer thresholds share with the human thresholds the elevation in the blue illumination-change direction relative to the red and green directions, indicating that the information available to the visual system at the retinal photoreceptor stage is already biased across illumination directions. However, the difference between the computational-observer thresholds between the blue and yellow illumination-change directions is very small and does not provide a clear explanation for the elevation of thresholds in the blue direction that is generally found in the human psychophysics, when thresholds are expressed using the CIELUV ΔE metric.

**Figure 6.**
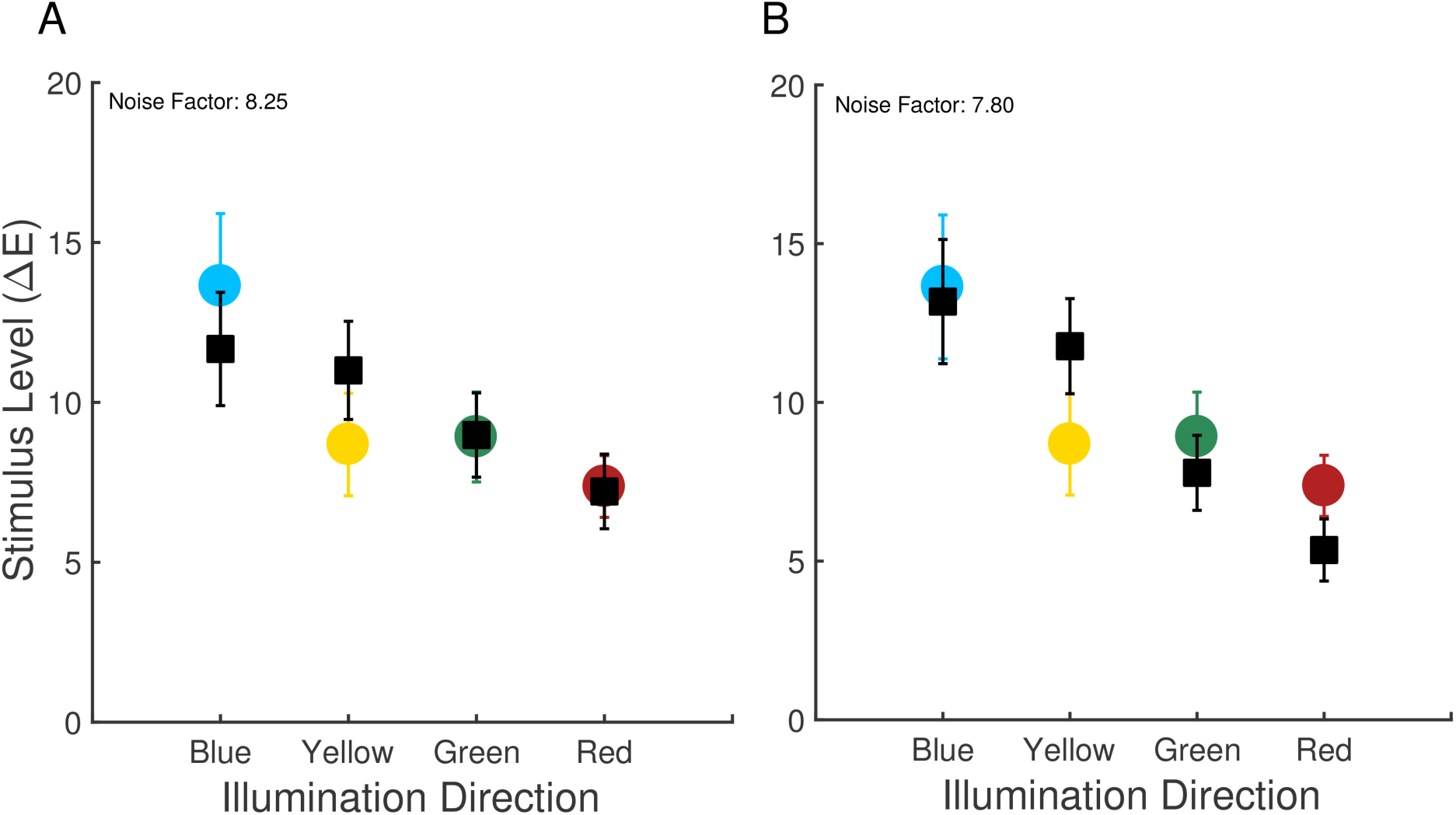
Comparison of computational and psychophysical performance: Experiment 1. **A)** Computational-observer thresholds (black squares) along with human observer thresholds. The noise factor leading to the best fit to each human observer’s data was found, and the resulting computational-observer thresholds were averaged over observers. Computational-observer thresholds were calculated using aggregation over all stimulus patches. Error bars are +/−1 SEM across observers. Symbol key is as in Figure 1. **B)** Same as A, except using computational-observer thresholds obtained with fixation-weighted aggregation.

The average (across observers) noise factor obtained in the fitting above was 8.25. This factor provides an omnibus summary of the net decrease in performance between the computational and human observers. There are many factors that likely contribute to the decrease, including neural noise introduced after the cone isomerizations, differences in spatial and temporal integration of information between the computational and human observers, loss of information during the inter-stimulus intervals, and differences in the decision processes used by the computational and human observers. The present work does not distinguish the contribution of each of these factors.

### Effect of eye fixations

The approach to aggregating the performance across patches described above weights the information from each stimulus patch equally. The fixations made by human observers during the experiment, however, show that they did not look at each part of the scene equally often. Figure 7A, 7B, and 7C illustrate the fixations made a by a single representative observer across both sessions of the experiment. Panel A shows the fixations made during the reference interval, while panels B and C show the fixations made during the first and second comparison intervals respectively. In all cases, the fixations generally cluster around one location in the scene, with the fixations made during the reference interval more spread out, presumably because the duration of the reference interval was longer (2370 ms vs. 870 ms). A more detailed analysis of the fixation data is provided in Radonjić et al. (2018).

**Figure 7.**
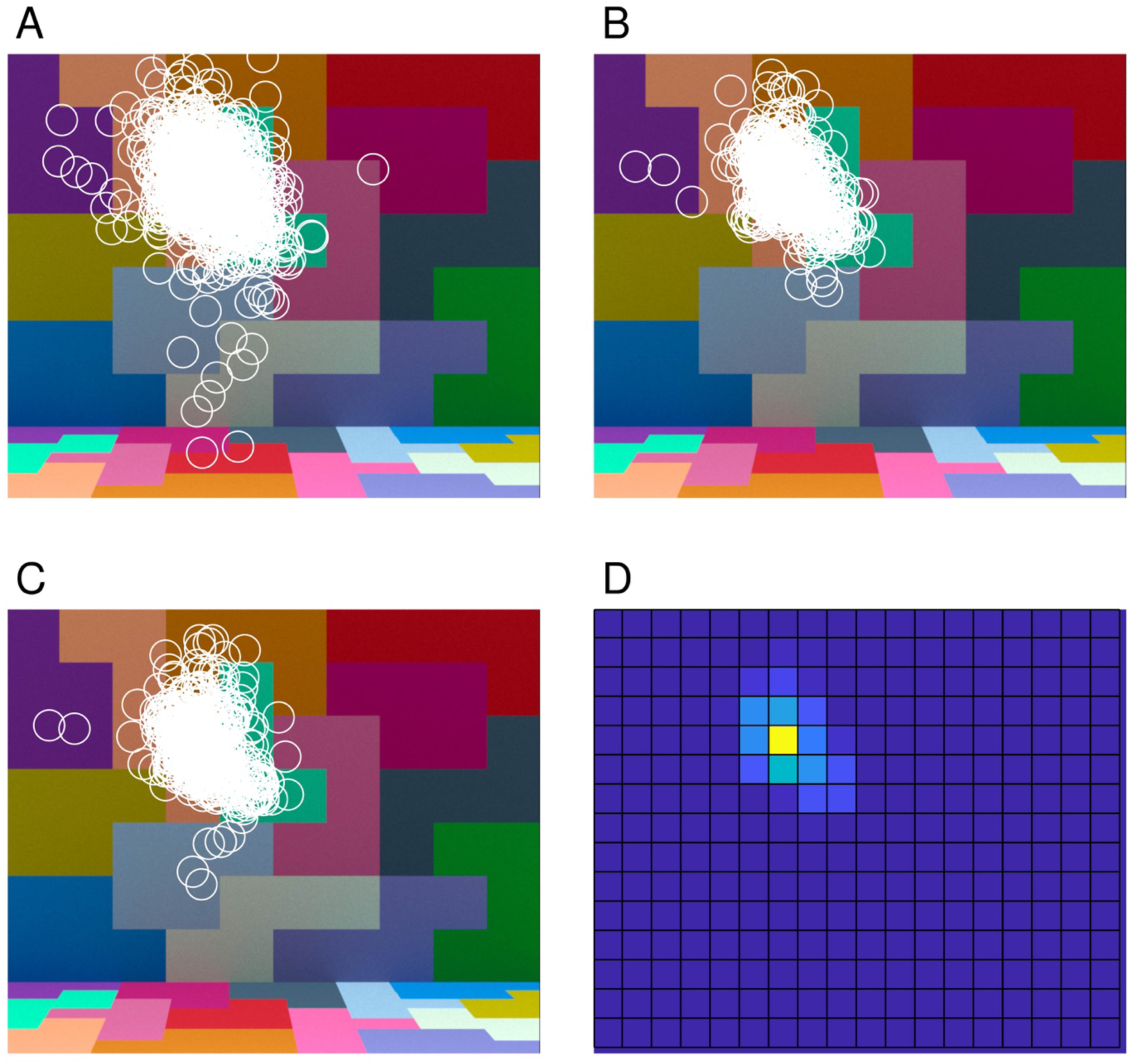
Eye fixation data. **A)** Fixations made by one observer (observer eom in Radonjić et al., 2018, note that observer initials are fictional and do not provide identifying information) during the reference interval, aggregated over all trials for that observer in both sessions. **B, C)** Fixations made by the same observer during the first and second comparison intervals. **D)** Distribution of combined fixations from B and C, represented as a heat map (yellow indicating a higher number of fixations and blue indicating a lower number). Each fixation was assigned to one of the 1° × 1° stimulus patches (see Figure 4A).

One approach to incorporating the fixation data into the computational-observer calculations is to weight each stimulus patch in the averaging step according to the fraction of fixations that landed within that patch during the two comparison intervals. Figure 7D illustrates the fixation-based weights for one observer as a heat map, with the weights for each patch obtained from the combined data shown in 7B and C. We incorporated fixation-based weights for each observer using that observer’s eye fixation data and the computed thresholds for each illumination-change direction/step size/noise factor. We then found the noise factor for each observer that brought the computational-observer data into best register with that observer’s data, and then averaged the thresholds across observers. The results are shown in Figure 6B. The effect of incorporating eye fixations is not large. The primary effect is the increase in the difference between the blue and yellow direction thresholds on the one hand the red and green direction thresholds on the other.

### Information provided by the different cone classes

The computational-observer thresholds are elevated in the blue and yellow directions relative to the red and green directions. This suggests that there is an asymmetry across illumination-change directions in the information available at the photoreceptor mosaic, when step size is expressed in ΔE units. We were curious about how the different cone classes contribute to this asymmetry. We recomputed computational-observer thresholds for six additional cone mosaics. These were mosaics with: L cones only, M cones only, S cones only, L and M cones, L and S cones, and M and S cones. In constructing the dichromatic cone mosaics, missing L cones were replaced with M cones and vice-versa, while missing S cones were replaced with a mixture of L and M cones in a 2:1 (L:M) ratio.

Figure 8 shows computational-observer thresholds for the six additional mosaics (black squares in each plot) along with the corresponding computational observer data for the original trichromatic mosaic (colored circles in each plot, replotted from black squares shown in Figure 6A). Computational-observer thresholds for the L, LM, and LS mosaics are very similar to one another as well to those obtained with the original mosaic. This suggests that the blue/yellow elevation in computational-observer thresholds seen in with the original mosaic can be primarily attributed to pattern of performance mediated by the L cones: mosaics containing L cones show the elevation, while those with only M and/or S cones do not. Prossibly, the fact that L cones are the predominate type in the mosaic leads to the overall mosaic showing a pattern of performance that is most similar to the performance of the L-cone-only mosaic. This in turn might mean that some of the individual variability observed in illumination-discrimination performance (see Radonjić et al., 2016; Radonjić et al., 2018) could be related to individual variability in L:M cone ratio (Hofer, Carroll, Neitz, Neitz, & Williams, 2005; for a review see Brainard, 2015). We note, however, that there are other possibilities. For example, there is a systematic difference in the isomerization rate between L and S cones because of light absorption by the lens and macular pigment. This difference might interact with our choice of SVM linear classifier or our choice to model post-receptoral processing with equal-variance Gaussian noise.

**Figure 8.**
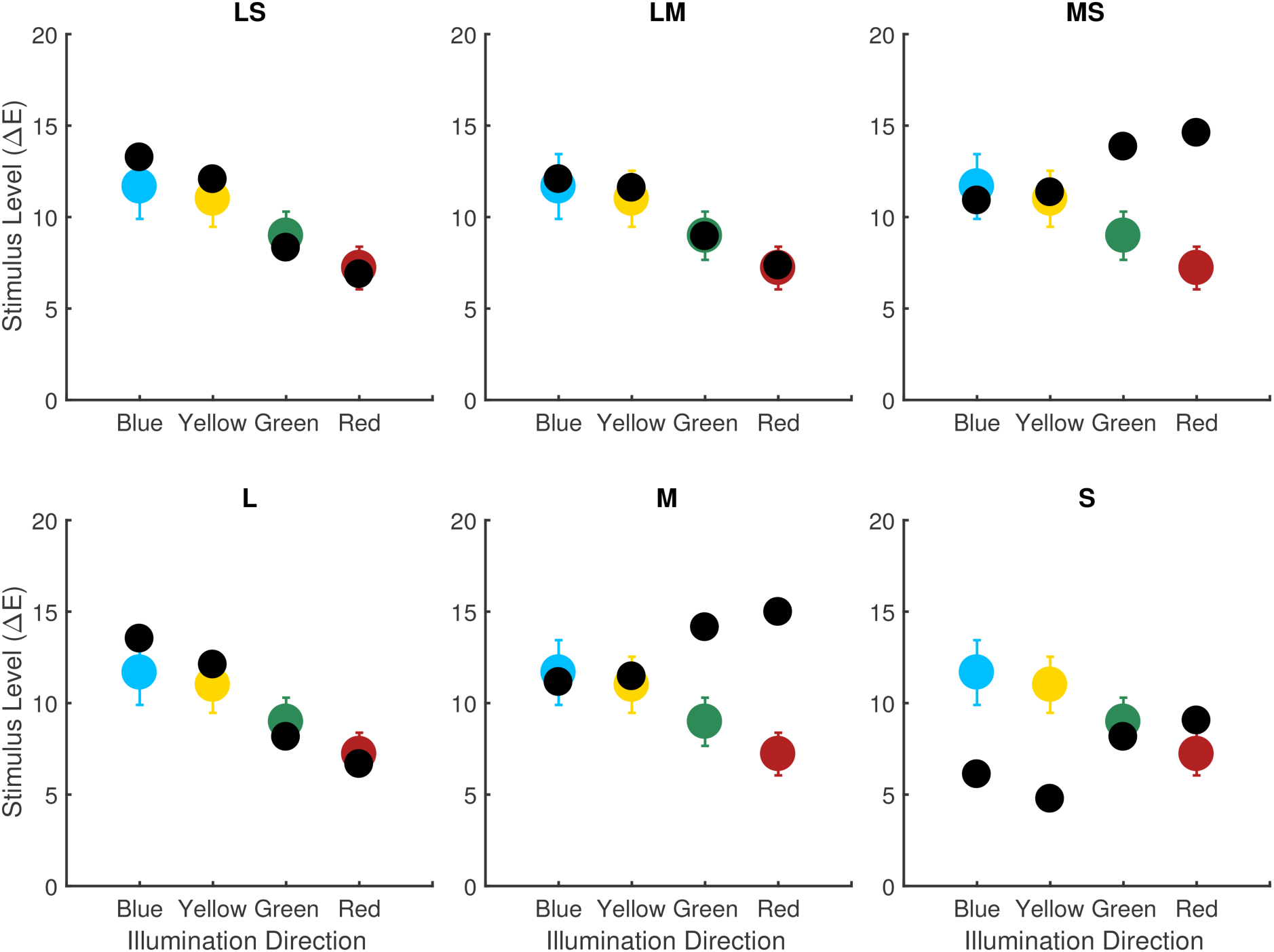
Effect of variation in mosaic. Best fit computational-observer thresholds for alternate mosaics (black squares in each panel) compared to the computational-observer thresholds for the trichromatic mosaic (colored circles in each panel, replotted from the black squares in Figure 6A). From left to right top row: LM cones only (no S, tritanope), LS cones only (no M, deuteranope), MS cones only (no L, protanope). From left to right bottom row: L cone only, M cone only, S cone only. Performance aggregation is over all stimulus patches, since we do not have fixation data corresponding to human observers with the alternate mosaics. The same noise factor (8.25) is used for all of the di- and monochromatic mosaic computational-observer thresholds shown here. This value is the average noise factor value obtained for the trichromatic computational-observer thresholds in 6A.

The relative computational-observer thresholds for M, S, and MS cone mosaics are also similar to one another, although sensitivity is lower overall for S, and these thresholds have a different pattern across illumination-change directions than that obtained with L-cone dominated mosaics. Here, computational-observer thresholds for the blue and yellow illumination directions are the lowest relative to the other color directions.

Aston et al. (2016) measured illumination discrimination thresholds in dichromats using stimuli similar to those we analyze. For protanopes (corresponding to our MS mosaic) there was a relative elevation of human thresholds in the red and green illumination-change directions, similar to what we see for the computational-observer thresholds. For deuteranopes (corresponding to our LS mosaic), however, thresholds in the red and green illumination-change directions were also elevated, which is inconsistent with the effect seen with the computational observer.

Alvaro et al. (2017) also measured illumination-discrimination threshold in dichromats, but using stimuli derived from natural hyperspectral images and only along blue and yellow illuminant-change directions. They found that thresholds were slightly elevated overall for dichromats relative to trichromats, with no statistically significant interaction between observer type and illuminant-change direction. Given that the changes in relative computational-observer thresholds across LMS, LS and MS mosaics for the blue and yellow directions are small, the lack of interaction may be consistent with the computational observer results.

### Modeling Illumination Discrimination: Modeled Experiment 2

Radonjić et al. (2016) reported results from an illumination discrimination experiment similar to the one described above, but where across conditions the ensemble of surfaces in the stimulus scene was varied. They obtained illumination discrimination thresholds for three separate scene surface ensembles, which they labeled neutral, yellowish-green and reddish-blue (based on the appearance of the corresponding images). The basic methodology of their experiment was the same as for Experiment 1 above, except for the variation of surface ensemble and a slightly different geometric structure of the stimulus scene. In addition, eye fixations were not measured. The detailed methods are provided in the published report (Radonjić et al., 2016). Figure 9 shows the illumination discrimination thresholds they measured for each of their surface ensemble conditions (panels D-F). Illumination discrimination thresholds depend strongly on the ensemble of surfaces in the scene. Radonjić et al. conjectured that the threshold variation was driven by changes in the information available in the image to make the discriminations. We investigate that conjecture here.

We computed computational-observer performance using the methods described above for the experimental stimuli of Radonjić et al. (Radonjić et al., 2016). Figure 9 shows the results of the analysis. The top row (panels A-C) shows the computational-observer thresholds as a function of the noise factor for the neutral, reddish-blue, and yellowish-green scene conditions. The bottom row (panels D-F) show the fits of the computational-observer to the psychophysical data. As above, the computational-observer noise factor was fit separately for each observer before averaging across observers. A single noise factor was found for each observer, held fixed across the three surface ensembles and four illumination-change directions. Since eye fixation data were not available, performance was aggregated over all stimulus patches with equal weight.

**Figure 9.**
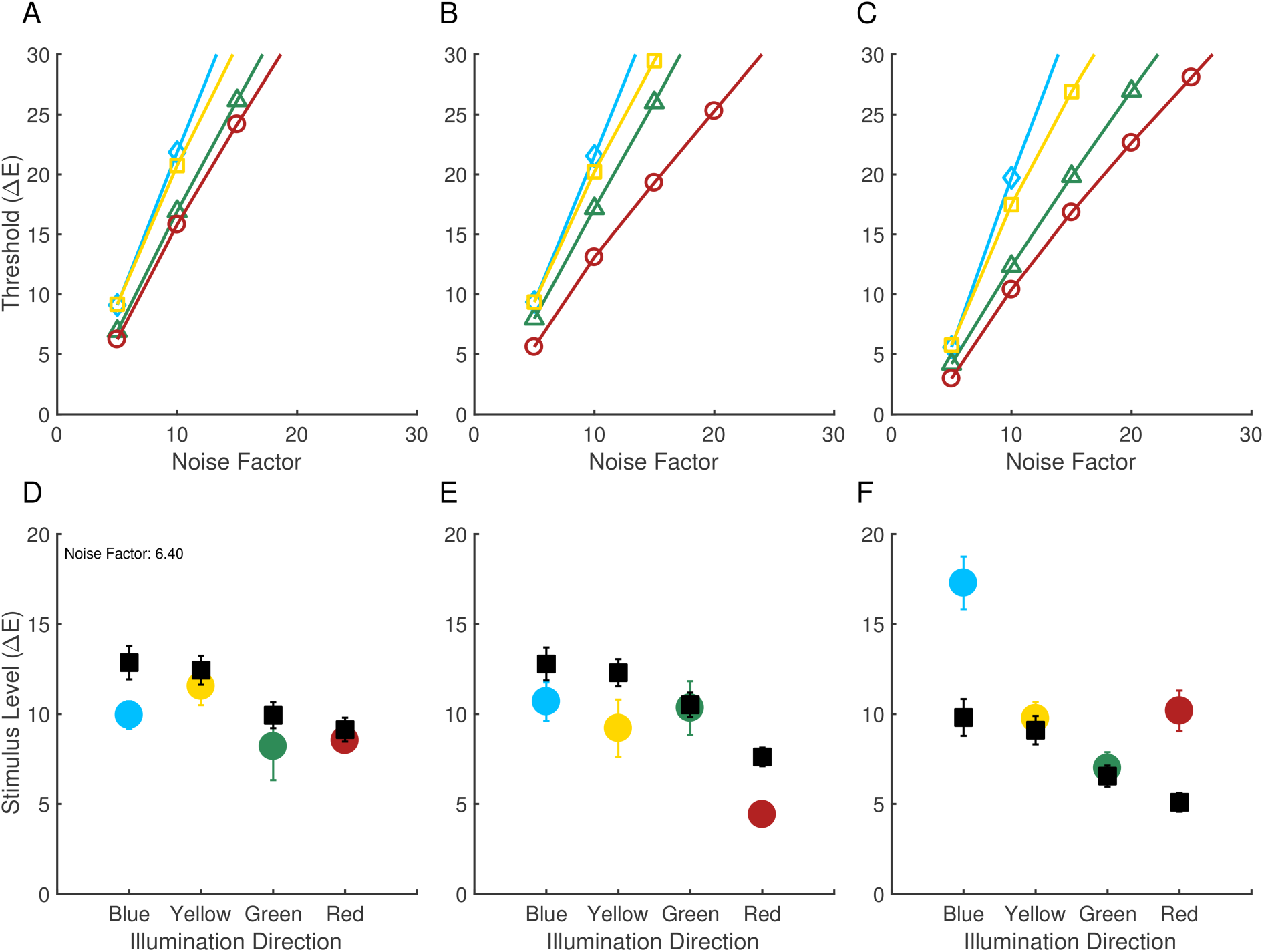
Comparison of computational and psychophysical performance: Experiment 2. Computational-observer thresholds as a function of noise factor for the neutral (panel A), reddish-blue (B) and yellowish-green condition (C). Computational-observer thresholds fit to human experimental data are also shown for the neutral (panels D), reddish-blue (E) and yellowish-green condition (F). Computational-observer performance was aggregated over all stimulus patches (without applying fixation-based weighting, since we do not have fixation data for these measurements). Fits for panels D-F share the same noise factor (6.4). The data in the neutral condition (panel A) represent an unusual case where human thresholds were not highest in the blue illumination-change direction.

Contrary to the conjecture of Radonjić et al. (2016), there is little variation in the computational-observer thresholds across the change in surface ensemble. The small effects seen are an overall decrease in predicted thresholds for the yellowish-green ensemble (panel F) and a relative decrease in threshold for the red illuminant-change direction for the reddish-blue ensemble (panel E). These two changes do not capture the two striking effects in the psychophysical data, a large decrease in red illuminant-change direction threshold between the neutral/yellowish-green and the reddish-blue ensemble, and a large increase in the blue illuminant-change direction threshold between the neutral/reddish-blue and the yellowish-green ensemble.

## Discussion

We implemented a computational observer that performs the illumination discrimination task. The computational observer has access to the representation at the cone mosaic and learns a linear SVM classifier. Overall, across the stimulus conditions we studied, thresholds determined from the computational observer’s performance do not predict the relative performance of human observers across illumination-change directions and across variation in the ensemble of surfaces in the scene. The deviations are particularly clear for the latter case, where we compared the computational observer to data reported by Radonjić et al. (2016). Whereas human performance showed a large effect of the surface ensemble, the computational observer’s performance stayed relatively consistent across the ensembles. The computational observer results are also inconsistent with Aston et aľ.s (2016) measurements of illumination-discrimination thresholds for deuteranopes.

### Limitations

Our computational observer incorporates a number of simplifications. One of these is to implement the computational observer on 1° × 1° patches evenly spaced across the stimulus, instead of simulating a larger mosaic with varying cone density that samples the entire stimulus. This choice was made for reasons of computational efficiency, particularly with respect to learning the classifier. As computers get faster, we will be able to move towards simulating larger and more realistic cone mosaics (with hexagonal packing and with cone density varying with eccentricity).

For Experiment 1, we compared computational-observer thresholds when we weighted all stimulus patches equally and when we weighted them according to observers’ measured eye fixations. The two analyses yielded similar patterns of thresholds. We were not able to make this comparison for Experiment 2 nor for hypothetical observers with di- and monochromatic cone mosaics, because we do not have fixation data available for those cases. There are cases where which patch the computational observer is trained and evaluated on has a large effect on performance (Figure 5A, B). This suggests that the information about where observers look when they perform psychophysical tasks is useful for computational observer modeling.

Other simplifications of our approach were that we assumed that observers used the information from only a single patch across the duration of each trial, and that we restricted our modeling to two comparison intervals, while excluding the reference interval. Building computational-observer performance models that take into account the trial-by-trial sequence of eye fixation locations across the entire trial is an interesting extension for future research. As with increasing the size of the modeled cone mosaic, this extension would also require an increase in computational resources.

### Post-isomerization Processes

Although it is of fundamental interest to understand how the information available in the responses of the cone mosaic varies across experimental conditions within a psychophysical study, it should not be surprising that there are cases where the rest of the visual system plays a role in shaping performance. Indeed, two well-known processes that occur after photopigment isomerization are likely to influence performance on the illumination-discrimination task. First, adaption that begins with the conversion of isomerization rate to photocurrent within the cones has the potential to affect post-receptoral information (Hood & Finkelstein, 1986; Stockman & Brainard, 2010; Angueyra & Rieke, 2013) and might specifically be expected to play a role in performance across different sets of surface ensembles.

Second, signals from the three classes of cones are recombined by post-receptoral processing into luminance and cone-opponent channels (Shevell & Martin, 2017). To the extent that noise following this recombination limits discrimination, these processes can have a major effect on the relative sensitivity of the visual system to stimulus changes in different directions in color space (for a review see Stockman & Brainard, 2010). Indeed, it has been shown that cone-opponent processing plays an important role in shaping sensitivity to modulations in different color directions in psychophysical tasks that involve simple colored stimuli, such as spots or Gabor patches seen against a spatially uniform background (Wandell, 1995; Stockman & Brainard, 2010).

Our current implementation of the computational observer uses additive zero-mean Gaussian noise as a proxy for all of post-receptor vision. This noise does not model stimulus-specific effects. Rather, it was included because real observer thresholds are considerably higher than those of the computational observer, even when we consider only 1° by 1° patches of retina. We are eager to extend our computational-observer models to incorporate both the conversion of isomerization rate to photocurrent and the recombination of signals from different classes of cones by retinal ganglion cells, along the lines of sequential ideal observer analysis outlined by Geisler (1989). A computational observer that models multiple stages of neural information processing in the retina will likely provide a better account of the experimental data with less need for an omnibus parameter to reduce overall efficiency.

## Acknowledgments

Supported by NIH EY10016 (DHB) and Simons Foundation Collaboration on the Global Brain Grant 324759 (DHB and BAW). We thank Stacey Aston and Anya Hurlbert for useful discussions about the work.

## Footnotes

1 The value of 1 ΔE is a nominal step size. The actual step sizes varied across illuminant directions and modeled experiments. The actual step sizes were used in all analyses. See Radonjić et al. (2016; 2018) for details.

2 Stimulus images in Modeled Experiment 1 were viewed from 68.3 cm and subtended 20.0 by 16.7 degrees of visual angle. Images in Modeled Experiment 2 were viewed from 76.4 cm and subtended 18.6 by 17.3 degrees of visual angle. The corresponding scene specifications in ISETBio used viewing distances identical to those in the experiments, but different sizes (Experiment 1: 18.0° by 15.1°; Experiment 2: 10.8° by 10.0°).

3 Because rendering is a stochastic process, it is possible to train a classifier to distinguish between two separate renderings of the same scene, even though the differences are imperceptible to a human observer. In Modeled Experiment 1, we used 30 separate renderings of the target image and drew randomly from these on each psychophysical trial to generate each training/test set instance. Our modeling followed this same procedure. In Modeled Experiment 2 described below, only one target image was used in the psychophysics. In the modeling, we generated 7 versions of that same target image and drew randomly from these to generate each training/test set instance. The two draws for each trial (one for target interval and the other for one of the comparison intervals) were made without replacement.

## References

Alvaro, L., Linhares, J. M. M., Moreira, H., Lillo, J., & Nascimento, S. M. C. (2017). Robust colour constancy in red-green dichromats. PLoS One, 12(6), e0180310. https://www.ncbi.nlm.nih.gov/pubmed/28662218.

Angueyra, J. M., & Rieke F. (2013). Origin and effect of phototransduction noise in primate cone photoreceptors. Nature Neuroscience, 16(11), 1692–1700. <Go to ISI>://WOS:000326205700028 http://www.nature.com/neuro/iournal/v16/n11/pdf/nn.3534.pdf.

Aston, S., Turner, J., Le Couteur Bisson, T., Jordan, G., & Hurlbert, A. C. (2016). Better colour constancy or worse discrimination? Illumination discrimination in colour anomalous observers. Perception (Supplement), 45(2), 207.

Banks, M. S., Geisler, W. S., & Bennett, P. J. (1987). The physical limits of grating visibility. Vision Research, 27(11), l915–1924.

Barlow, H. B. (1962). Measurements of the quantum efficiency of discrimination in human scotopic vision. Journal of Physiology, 160(1), 169–188. http://ip.physoc.org/content/160/1/169.full.pdf.

Beck J. (1959). Stimulus correlates for the judged illumination of a surface. Journal of Experimental Psychology, 58(4), 267–274.

Bone, R. A., Landrum, J. T., & Cains A. (1992). Optical density spectra of the macular pigment in vivo and in vitro. Vision Research, 32(1), 105–110. http://ac.els-cdn.com/0042698992901183/1-s2.0-0042698992901183-main.pdf?tid=2dd6b810-da92-11e2-8479-00000aab0f01&acdnat=1371833394d159994bb8149718b27a187f653ee9db.

Brainard, D. H. (2015). Color and the cone mosaic. Annu Rev Vis Sci, 1, 519–546. https://www.ncbi.nlm.nih.gov/pubmed/28532367.

CIE. (2004). Colorimetry, third edition (No. 15.2004). Vienna: Bureau Central de la CIE.

CIE. (2007). Fundamental chromaticity diagram with physiological axes – Parts 1 and 2. Technical Report 170-1. Vienna: Central Bureau of the Commission Internationale de ľ Éclairage.

Curcio, C. A., Allen, K., Sloan, K., Lerea, C., Klock, I., & Milam A. (1991). Distribution and morphology of human cone photoreceptors stained with anti-blue opsin. Journal of Comparative Neurology, 312, 610–624.

Farrell, J. E., Jiang, H., Winawer, J., Brainard, D. H., & Wandell, B. A. (2014). Modeling visible differences: the computational observer model. Paper presented at Proceedings of the 2014 Society for Information Display (SID) International Symposium.

Geisler, W. S. (1989). Sequential ideal-observer analysis of visual discriminations. Psychological Review, 96(2), 267–314. http://graphics.tx.ovid.com/ovftpdfs/FPDDNCJCQCNCPG00/fs047/ovft/live/gv024/00006832/00006832-198904000-00005.pdf.

Geisler, W. S. (2011). Contributions of ideal observer theory to vision research. Vision Research, 51(7), 771–781. http://www.ncbi.nlm.nih.gov/pubmed/20920517 http://ac.els-cdn.com/S0042698910004724/1-s2.0-S0042698910004724-main.pdf?tid=8d26b4ca-d832-11e3-81c7-00000aab0f27&acdnat=1399719918a7f3815a4cf9c3ed2804c54cd1eb3d1c.

Gilchrist, A., & Jacobsen A. (1984). Perception of lightness and illumination in a world of one reflectance. Perception, 13, 5–19.

Green, D. M., & Swets, J. A. (1966). Signal detection theory and psychophysics. New York: Wiley.

Hecht, S., Schlaer, S., & Pirenne, M. H. (1942). Energy, quanta and vision. Journal of the Optical Society of America, 38(6), 196–208. http://www.ncbi.nlm.nih.gov/pubmed/19873316.

Hernandez-Andres, J., Romero, J., Nieves, J. L., & Lee, R. L. (2001). Color and spectral analysis of daylight in southern Europe. Journal of the Optical Society of America a-Optics Image Science and Vision, 18(6), 1325–1335. <Go to ISI>://WOS:000168938900014.

Hofer, H., Carroll, J., Neitz, J., Neitz, M., & Williams, D. R. (2005). Organization of the human trichromatic cone mosaic. Journal of Neuroscience, 25, 9669–9679.

Hood, D. C., & Finkelstein, M. A. (1986). Sensitivity to light. In K. Boff, L. Kaufman & J. Thomas (Eds.), Handbook of Perception and Human Performance (Vol. 1, pp. 5-1–5-66). New York: Wiley.

Hurlbert, A. (1989). Cues to the color of the illuminant. Unpublished PhD thesis, Massachusetts Institute of Technology, Cambridge, MA.

Jiang, H., Cottaris, N. P., Golden, J., Brainard, D., Farrell, J. E., & Wandell, B. A. (2017). Simulating retinal encoding: factors influencing Vernier acuity. Paper presented at Electronic Imaging 2017, Burlingame, CA.

Kardos L. (1928). Dingfarbenwahrnehmung und Duplizitätstheorie. Zeitschrift für Psychologie, 108, 240–314.

Kingdom, F. A. A., & Prins, N. (2010). Psychophysics: A Practical Introduction. San Diego, CA: Academic Press.

Kolb, H. Photoreceptors. Webvision Retrieved March 30, 2018, from http://webvision.med.utah.edu/book/part-ii-anatomv-and-phvsiology-of-the-retina/photoreceptors/

Kozaki, A., & Noguchi K. (1976). The relationship between perceived surface-lightness and perceived illumination. Psychological Research, 39(1–16)

Lee, T. Y., & Brainard, D. H. (2011). Detection of changes in luminance distributions. Journal of Vision, 10(13:14) https://www.ncbi.nlm.nih.gov/pubmed/22085597.

Logvinenko, A., & Menshikova G. (1994). Trade-off between achromatic colour and perceived illumination as revealed by the use of pseudoscopic inversion of apparent depth. Perception, 23(9), 1007–1023.

Logvinenko, A. D., & Maloney, L. T. (2006). The proximity structure of achromatic surface colors and the impossibility of asymmetric lightness matching. Perception & Psychophysics, 68(1), 76–83. http://www.ncbi.nlm.nih.gov/pubmed/16617831.

Lopez, H., Murray, H. L., & Goodenough, D. J. (1992). Objective analysis of ultrasound images by use of a computational observer. IEE Transactions on Medical Imaging, 11(4), 496–506. <Go to ISI>://WOS:A1992KK63500005.

Manning, C. D., Raghavean, P., & Schutze, H. (2008). Introduction to Information Retrieval. Cambridge: Cambridge University Press.

Marimont, D. H., & Wandell, B. A. (1994). Matching color Images - the effects of axial chromatic aberration. Journal of the Optical Society of America A, 11(12), 3113–3122. <Go to ISI>://WOS:A1994PX50900001.

Nascimento, S. M., Amano, K., & Foster, D. H. (2016). Spatial distributions of local illumination color in natural scenes. Vision Res, 120, 39–44. http://www.ncbi.nlm.nih.gov/pubmed/26291072.

Noguchi, K., & Kozaki A. (1985). Perceptual scission of surface lightness and illumination: An examination of the Gelb effect. Psychological Research, 47(1), 19–25.

Oyama T. (1968). Stimulus determinants of brightness constancy and the perception of illumination. Japanese Psychological Research, 10(3), 146–155.

Pearce, B., Crichton, S., Mackiewicz, M., Finlayson, G. D., & Hurlbert A. (2014). Chromatic illumination discrimination ability reveals that human colour constancy is optimised for blue daylight illuminations. PLoS ONE 9(2:e87989), e87989. http://www.ncbi.nlm.nih.gov/pubmed/24586299.

Radonjić, A., Ding, X., Krieger, A., Aston, S., Hurlbert, A. C., & Brainard, D. H. (2018). Illumination discrimination in the absence of a fixed surface-reflectance layout. In press

Radonjić, A., Pearce, B., Aston, S., Krieger, A., Dubin, H., Cottaris, N. P., Brainard, D. H., & Hurlbert, A. C. (2016). Illumination discrimination in real and simulated scenes. Journal of Vision, 16(11:2), 1–18. http://dx.doi.org/10.1167/16.1L2 http://iov.arvoiournals.org/data/iournals/iov/935705/i1534-7362-16-11-2.pdf.

Rodieck, R. W. (1998). The First Steps in Seeing. Sunderland, Mass.: Sinauer.

Rutherford, M. D., & Brainard, D. H. (2002). Lightness constancy: a direct test of the illumination estimation hypothesis. Psychological Science, 13, 142–149.

Sekiguchi, N., Williams, D. R., & Brainard, D. H. (1993). Efficiency in detection of isoluminant and isochromatic interference fringes. Journal of the Optical Society of America A, 10(10), 2118–2133. http://www.opticsinfobase.org/DirectPDFAccess/38814A9A-AEF3-EBE9-5EF88C2F072A447A4820/iosaa-10-10-2118.pdf?da=1&id=4820&seq=0&mobile=no.

Shevell, S. K., & Martin, P. R. (2017). Color opponency: tutorial. J Opt Soc Am A Opt Image Sci Vis, 34(7), 1099–1108. https://www.ncbi.nlm.nih.gov/pubmed/29036118.

Spitschan, M., Aguirre, G. K., Brainard, D. H., & Sweeney, A. M. (2016). Variation of outdoor illumination as a function of solar elevation and light pollution. Scientific Reports, 6, 26756.

Stockman, A., & Brainard, D. H. (2010). Color vision mechanisms. In M. Bass, C. DeCusatis, J. Enoch, V. Lakshminarayanan, G. Li, C. Macdonald, V. Mahajan & E. van Stryland (Eds.), The Optical Society of America Handbook of Optics, 3rd edition, Volume III: Vision and Vision Optics (pp. 11.11–11.104). New York: McGraw Hill.

Stockman, A., Sharpe, L. T., & Fach, C. C. (1999). The spectral sensitivity of the human short-wavelength cones. Vision Research, 39(17), 2901–2927. http://ac.els-cdn.com/S0042698998002259/1-s2.0-S0042698998002259-main.pdf?tid=69561354-da88-11e2-a09a-00000aacb360&acdnat=13718291998998a5e4e157b73707f7369b7fd3a9ce.

Wandell, B. A. (1995). Foundations of Vision. Sunderland, MA: Sinauer.

Wetherill, G. B., & Levitt H. (1965). Sequential estimation of points on a psychometric function. The British Journal of Mathematical and Statistical Psychology, 18, 1–10.

Wyszecki, G., & Stiles, W. S. (1982). Color Science: Concepts and Methods, Quantitative Data and Formulae (2nd ed.). New York: John Wiley & Sons.

